# Slow conformational changes in the rigid and highly stable chymotrypsin inhibitor 2

**DOI:** 10.1101/2022.12.21.521530

**Authors:** Yulian Gavrilov, Andreas Prestel, Kresten Lindorff-Larsen, Kaare Teilum

**Author notes:** Corresponding author: Kaare Teilum.

## Abstract

Slow conformational changes are often directly linked to protein function. It is however less clear how such processes may perturb the overall folding stability of a protein. We previously found that the stabilizing double mutant L49I/I57V in the small protein chymotrypsin inhibitor 2 from barley led to distributed increased nano second and faster dynamics. Here we asked what effect this mutant and the two individual mutants L49I and I57V have on the slow conformational dynamics of CI2. We used ^15^N CPMG spin relaxation dispersion experiments to measure the kinetics, thermodynamics and structural changes associated with slow conformational change in CI2. These changes result in an excited state that is populated to 4.3% at 1 °C. As the temperature is increased the population of the excited state decreases. Structural changes in the transition to the excited state are associated with residues that interact with water molecules that have well defined positions and are found at these positions in all crystal structures of CI2. The mutations in CI2 have only little effect on the structure of the excited state whereas the stability of the excited state to some extent follows the stability of the main state. The minor state is thus most populated for the most stable CI2 variant and least populated for the least stable variant. We hypothesize that the interactions between the mutated residues and the well-ordered water molecules links subtle structural changes around the mutated residues to the region in the protein that experience slow conformational changes.

## Introduction

CI2 is a small single domain protein of 64 residues that is a highly efficient inhibitor of fungal subtilisins^1^ and has been extensively used as a model to understand key concepts of protein folding and stability^2–6^. The folding is following a highly cooperative two-state mechanism where the structure in the transition state forms around a partly formed structural nucleus in what is described as a nucleation condensation process^5^. Thus, only the folded native state and the disordered unfolded state have been observed in CI2 with some heterogeneity in the unfolded state due to proline isomerization^3^.

CI2 is highly stable with a free energy for folding, Δ*G*_f_ = 31 kJ/mol at 25 °C, and heat unfolds at 79 °C^5,7^. Single point mutations generally destabilize the structure, except for substitutions at positions 48 and 55 that may lead to significant stabilization^8,9^. We recently demonstrated that a set of point mutations (L49I and I57V) synergistically stabilize the native state of chymotrypsin inhibitor 2, CI2, by 5.1 kJ/mol more than the effect of the individual mutations^9^. We also showed that the increase in stability correlates with increased conformational entropy distributed throughout the molecule^10^.

Molecular motions in proteins occur on timescales ranging from femtoseconds to picoseconds for atomic vibrations, and up to hours for rearrangements of subunits and for folding of some proteins^11^. The fast molecular motions on the nano-second timescale and faster contribute to the conformational entropy of the molecule, whereas slower motions are associated with distinct conformational states that may be directly coupled to protein function^12,13^. In a few proteins, motions on the μs-ms time scales have been coupled to conformational changes around long-lived water molecules bound to the protein^14,15^. Nuclear spin relaxation measured by NMR spectroscopy allow the quantification at atomic resolution of motions on both the sub-ns time scale and on the time scales from μs to sec^16^.

Our analysis of the fast sub-nanosecond dynamics revealed that CI2 also experiences conformational dynamics on the ms time scale, but from that work we could not extract any details about the slow process^10^. In the present work, we asked what the characteristics of the slow conformational process are and if this might help rationalize some of the stabilizing effect in the L49I/I57V variant of CI2. We have used ^15^N CPMG spin relaxation dispersion experiments to measure the kinetics, thermodynamics and structural changes associated with the conformational change^17^. Spin relaxation dispersion experiments are widely used to characterize millisecond conformational exchange, and in many proteins structural movements associated with protein function have been identified^18^.

We find substantial slow dynamics in CI2, mostly localized to one region of the protein. Specifically, we find that millisecond conformational changes in CI2 are associated with residues that interact with water molecules that are well defined in crystal structures of CI2. These motions are preserved in all four variants of CI2 that we studied (wild-type, L49I, I57V and L49I/I57V), though with subtle differences that to some extent correlate with the overall stability of the proteins.

## Results

We measured ^15^N-CPMG relaxation dispersion profiles on WT CI2 at six different temperatures ranging from 1 °C to 25 °C and observed several residues that at all temperatures show measurable exchange contributions to the transverse relaxation, while other residues have completely flat profiles (Figure 1A and B, Figure S1-S6). The magnitude of the exchange contribution, *R*_ex_, clearly decreases with temperature (Figure 1A). Thus, the number of residues that meet our selection criteria for further analysis (*R*_ex_ > 2 s^-1^) changes from five at 25 °C to 20 at 1 °C (Table 1). For each dataset, we fitted all selected residues globally to a two-state model (Figure S1-S6):

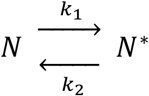

yielding the rate for the exchange between the two states, *k*_ex_ = *k*_1_ + *k*_2_, the population of the minor state, *p*_N*_, and for each residue the chemical shift difference between the exchanging states, |Δω| = |ω_N*_ - ω_N_|. For the data at 25 °C, *k*_ex_ is very large, which results in *p*_N*_ and |Δ*ω*| becoming highly correlated ^19^. Consequently, at 25 °C it is only possible to extract ϕ = *p*_N*_(1-*p*_N*_)Δω^2^ in addition to *k*_ex_.

**Figure 1.**
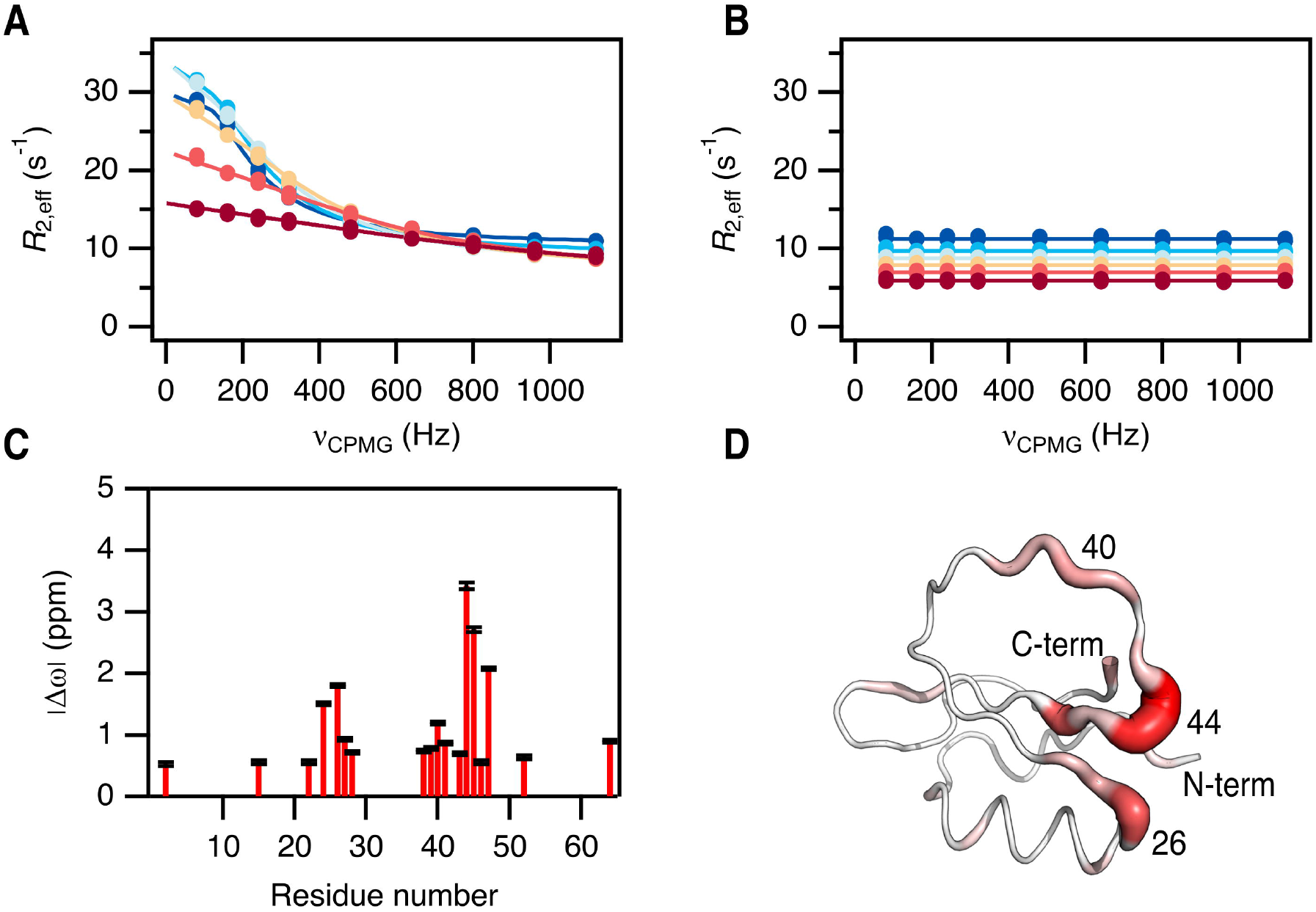
Conformational exchange in CI2 from ^15^N Relaxation dispersion of backbone amides. Representative dispersions (*R*_2,app_ vs ν_CPMG_) at a static field of 75 MHz for **A)** I44 that show large dispersions and **B)** K11 that shows no dispersion. The dispersions were recorded at six different temperatures (1 °C, 5 °C, 10 °C, 15 °C, 20 °C and 25 °C ranging from dark blue over yellow to dark red). Solid lines represent global fits to a two-state exchange model. **C)** Chemical shift change (|Δω|) from the global fit at 5 °C as a function of residue number. **D)** Structure of CI2 (PDB id 7A1H). The width of the backbone trace is scaled with |Δω|. The color scale is from white (0 ppm) to red (4 ppm).

**Table 1.** Overview of number of residues included in and the results of fitting relaxation dispersion data.

| Variant | Temp (°C) | n<sub>res</sub> | $k_{\text{ex}}$ (s<sup>-1</sup>) | $p_{\text{N}^*}$ (%) |
| --- | --- | --- | --- | --- |
| WT | 1 | 20 | $(52 \pm 1) \times 10$ | $4.3 \pm 0.1$ |
| WT | 5 | 18 | $(78 \pm 2) \times 10$ | $4.2 \pm 0.1$ |
| WT | 10 | 14 | $(130 \pm 3) \times 10$ | $3.9 \pm 0.1$ |
| WT | 15 | 11 | $(195 \pm 7) \times 10$ | $2.4 \pm 0.2$ |
| WT | 20 | 6 | $(31 \pm 2) \times 10^2$ | $2.2 \pm 0.6$ |
| WT | 25 | 5 | $(58 \pm 5) \times 10^2$ | NA<sup>a</sup> |
| L49I | 5 | 15 | $(76 \pm 3) \times 10$ | $3.6 \pm 0.1$ |
| I57V | 5 | 17 | $(83 \pm 2) \times 10$ | $4.0 \pm 0.1$ |
| L49I/I57V | 5 | 18 | $(75 \pm 2) \times 10$ | $4.8 \pm 0.1$ |

<sup>a</sup>Data fit to a model where $p_{\text{N}^*}$ is not an individual fitting parameter.

The residues that experience the largest chemical shift difference between the exchanging states cluster in two regions around residue 25 and around residue 45 as demonstrated for the data at 5 °C (Figure 1C). Plotting the chemical shift changes on the structure of CI2 shows that these regions are close in space at one end of the central β-strands (Figure 1D). Also, there are smaller exchange contributions on the C-terminus and residues 38-41 in the middle of the inhibitory loop.

As we increased the temperature from 1 °C to 20 °C, *p*_N*_ decreased from 4.3% to 2.2% (Table 1). We have plotted the corresponding Δ*G* values against the temperature in Figure 2A and observe a non-linear temperature dependence. We fitted the data to Eq (1) to find the enthalpy, Δ*H* = 7.05 ± 0.05 kJ/mol, for the reaction at the temperature, *T*_0_ = −1 ± 2 °C, where the minor state is most stable relative to the major state. This temperature is where Δ*S* for the reaction is 0. The fit also gave Δ*C*_p_ = −2.2 ± 0.6 kJ/mol/K. As the temperature is increased the minor state thus becomes more enthalpically favorable, but also much more entropically unfavorable resulting in the decreasing *p*_N*_. We note that Δ*C*_p_ is negative in contrast to Δ*C*_p_ for protein unfolding, which is positive as a result of the larger exposed hydrophobic surface area of the unfolded state. The magnitude of the Δ*C*_p_ value for the N-to-N* transition (−2.2 kJ/mol/K) may, for example, be compared to that of full unfolding of CI2 (4.3 kJ/mol/K)^7^, demonstrating that the value is relatively large. The common interpretation of a negative Δ*C*_p_ value suggests that the transition results in a decrease in solvent accessible surface area.

**Figure 2.**
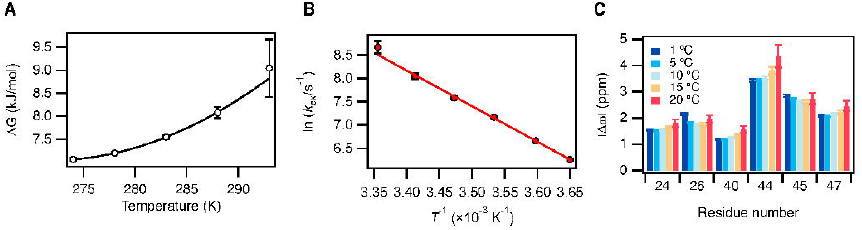
Temperature dependence of parameters extracted from global fits to a two-state model. **A)** Δ*G* calculated from the fitted *p*_N*_ as a function of temperature. The solid line is a fit to Eq. 1. **B)** Eyring plot of the fitted exchange rate, *k*_ex_. The fitted straight line has a slope of Δ*H*^‡^ = 63 kJ/mol. **C)** Chemical shift differences for the six residues that show significant dispersion at all temperatures from 1 °C to 20 °C.

We observed a large change in *k*_ex_ as the temperature was increased (Table 1). A linear regression of the Eyring plot of the measured rates at all temperatures results in an activation enthalpy of Δ*H*^‡^ = 63 kJ/mol (Figure 2B). As *p*_N*_ ≪ *p*_N_, *k*_ex_ is dominated by *k*_2_ and the Eyring plot thus gives the enthalpy of activation for the transition from the minor to the major state. This value is of the same order of magnitude as the activation enthalpies determined for the major folding phase of CI2^3^.

The chemical shift difference between the major and the minor states, |Δω|, show only minor changes with temperature for the six residues we can compare at all temperatures from 1 °C to 20 °C (Figure 2C). Most of the residues have increasing |Δω|, except Asp-45, for which |Δω| decreases slightly with temperature. Although the differences are subtle, we speculate that the different temperature coefficients for the major and minor states could be a result of changes in the hydrogen bonding in the two states.

In addition to WT CI2, we also analyzed three variants L49I, I57V and L49I/I57V. We again measured ^15^N-CPMG relaxation dispersions of the three variants, although only at 5 °C (Figure 3A and S7-9). The variants behave like the WT. However, we observe a variation in the dispersion step (*R*_ex_) among the variants, as exemplified for residue I44 (Figure 3A). From the fits of the experimental data, we see that *p*_N*_ varies with the amplitude of the dispersion step (Table 1). In contrast, there is little variation in |Δω| for the 12 residues which we can compare for all four variants (Figure 3B). The variation in *k*_ex_ is small ranging from 750 ± 20 s^-1^ to 830 ± 20 s^-1^ (Table 1).

**Figure 3.**
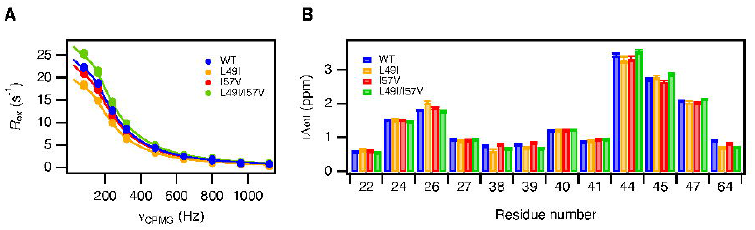
Conformational exchange for four CI2 variants. **A)** Dispersions for I44 at a static field of 75 MHz and 5 °C. **B)** Chemical shift differences for the set of 12 residues that shows dispersions in all four variants at 5 °C. Solid lines represent global fits to a two-state exchange model.

## Discussion

Our ^15^N-CPMG relaxation dispersion experiments demonstrate that CI2 undergoes slow conformational changes on the milli-second timescale. The most affected regions are clustered at the N-terminal end of the two main β-strands and in the inhibitory loop. These areas of CI2 already stood out as exchange outliers in our recent analysis of the fast dynamics in CI2^10^. The present analysis provides more details about the thermodynamics, and kinetics of this slow process, as well as the structures of the involved states. *p*_N*_ decreases with temperature and from the thermodynamic parameters extracted from our fits we estimate *p*_N*_ = 1.9% at 25 °C, which is the temperature at which most experimental analysis of the folding process of CI2 have been carried out^2,5,20^. At 25 °C, *k*_ex_ is more that 5000 s^-1^ and consequently the conformational exchange between the major and minor conformations of folded CI2 will be in rapid equilibrium compared with the folding process. From our data we cannot determine if both the major and the minor states can unfold directly. In any case, it is highly unlikely that the presences of the minor state will affect any conclusions about the two-state folding behavior of CI2^2,5,20^.

Based on the thermodynamic parameters for the N to N* process that we get from our present work, we find that *p*_N*_ is maximal at −1 °C and levels off at higher temperatures and becomes less than 0.1% at 54 °C (Figure 4). If we compare with how the population of the folded state, *p*_N_, decreases with the temperature, we see that N* is not present at any measurable level when the temperature unfolding begins (Figure 4). This observation strongly suggests that N* is not an intermediate obligatory for unfolding of CI2, which is also supported by the two-state behavior of CI2 folding.

**Figure 4.**
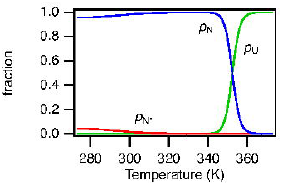
Variation with temperature of the population of the major, *p*_N_, and minor, *p*_N*_, folded states. The unfolded state, which is not shown, starts to dominate above 350 K.

The overall folding stability of the four analyzed variants of CI2 ^9^ to some extent follows how populated N* is at 5 °C (Table 1). Thus, *p*_N*_ is most populated for the most stable L49I/I57V variant (4.8%) and least populated for the least stable variant L49I (3.6%). The minor state of the WT and I57V are almost equally populated at 4.2% and 4.0%, respectively. However, the folding stability of I57V is slightly higher than that of WT.

The variation in *p*_N*_ is indeed significant and the difference between the highest and lowest value that we find for the variants at 5 °C, corresponds to the effect of changing the temperature more than 10 °C for the WT.

The existence of a high-energy native-like state, populated by around 4% of the folded CI2 molecules, will on its own only have a minor effect on the observed stability of the protein. We know from our previous work that the stability differences of the CI2 variants that we have studied here are to a large extent a result of differences in conformational entropy that are distributed throughout the molecule. The structures of the excited states of all four CI2 variants are very similar to each other as the chemical shift differences to the ground state are more or less identical (Figure 3B). In addition, positions 49 and 57 that are mutated are found in the hydrophobic core at the opposite end of the β-sheet relative to where we see the largest conformational changes. To summarize, the mutations at position 49 and 57 influence the relative stability of the minor state without perturbing the structure of this state.

From the relaxation dispersion data, we know which parts of CI2 that undergo the largest conformational changes, but with only chemical shift differences for the backbone amide ^15^N we do not have enough information to calculate a structure of the N* state. Previously, it has been shown that variation across crystal structures of a protein can help shed light on its dynamics in solution^21^. Thus, we speculated that we might get some hints about how the structure of N* looks by comparing the conformational variation of the structures of CI2 available from the PDB. Using the bio3D package in R^22^, we made a blast search in PDB and found 31 protein structures with sequences highly similar (−log(Evalue) > 84) to the sequence in the NMR structure 3CI2. Except from 3CI2 all structures are determined by X-ray crystallography. 19 are of CI2 in complex with subtilases and 12 are of CI2 alone. One structure, 6QIZ, is of a domain-swapped hexameric form of CI2^23^. As this is very different from the other structures, we excluded this from our analysis. We aligned the structures of the remaining 30 structures (Table S1) and calculated the root-mean-square-fluctuation of the C^α^ atoms (Figure 5A). The fluctuations are in general small and the largest fluctuations in the ensemble of structures are for positions 35-45 with the maximum observed for Met-40. This region makes up the inhibitory loop that is not stabilized by a hydrophobic core but rather held together by a large network of hydrogen bonds^24^. The region coincides with the residues that had slightly suppressed backbone order parameters in our previous NMR relaxation analysis^10^, which suggests CI2 being slightly more dynamic in the loop than in the rest of the protein on the nano-second timescale. The region where we see the largest chemical shift changes from the relaxation dispersions and thus has the largest change of the environment around the amide groups on the milli-second timescale is slightly shifted compared to the fast dynamics region and covers positions 38-47.

**Figure 5.**
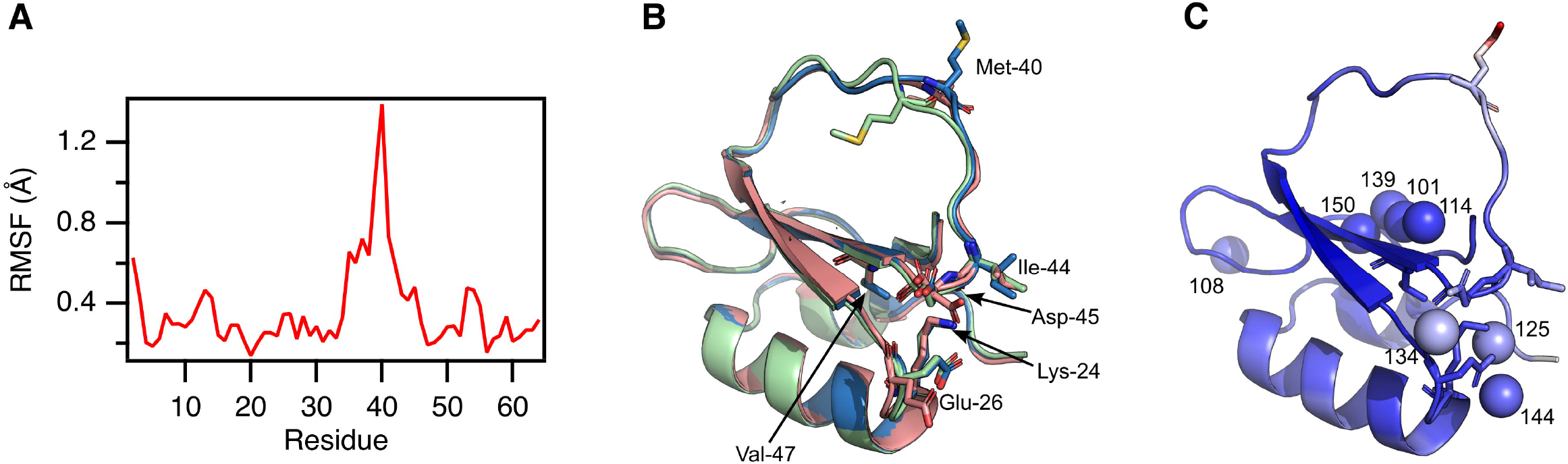
Structural variation in 30 experimental structures of CI2. **A)** Residue specific rootmean-square-fluctuation, RMSF. **B)** Three representative structures demonstrating the variation in the inhibitory loop and the side chain of Met-40. The side-chains of the six residues with significant dispersion at all temperatures from 1 °C to 20 °C are show in sticks. **C)** The eight water molecules with preserved positions in crystal structures are shown as spheres. The index of the water molecules in PDB entry 7a3m (CI2, I57V) are show. The same side-chains as in panel B are shown in sticks. The structure is colored according to the crystallographic B-factor in PDB entry 7a3m which ranges from 6 Å^2^ blue to 40 Å^2^ red.

We manually compared the structures of CI2 and found that the largest difference in side-chain fluctuations is seen for Met-40 and its position falls in three major classes. Most structures of free CI2 have the side chain of Met-40 pointing orthogonal to the inhibitory loop and away from the rest of the structure (blue structure in Figure 5B). In three of the structures of free CI2, Met-40 points towards the central β-strand and Arg-48 (green structure in Figure 5B). In the NMR structure of free CI2 and the structures of CI2 in complex with subtilisin, Met-40 also points towards the core of the protein (red structure in Figure 5B), but on the other side of the inhibitory loop compared with the green structure. The position of Met-40 appears to be connected to subtle structural changes in the side-chain positions of residues 28, 37-41, 43 and 48. Notably, these residues except Met-40 do not experience slow exchange of their backbone N^H^, and all side-chains of the residues that do experience exchange on their backbone N^H^ (again except Met-40) have very little change in the positions of their side-chains in the different structures.

In all the crystal structures of free and subtilase bound CI2, WT as well as mutants, six well defined water molecules appear at the same specific positions (Figure 5C). In 12 of the structures one additional water molecule, and in 11 structures two additional water molecules appear, corresponding waters number 125 and 144 in PDB entry 7a3m of CI2, I57V (Table S1). Four of the bound water molecules (101, 114, 138 and 150) lie partly buried in a chain along the central β-strand, which was also pointed out previously^25,26^. The most buried of these (number 150) is situated in the hydrophobic core of the protein between the side-chains of residue 49 and 57. The crystallographic B-factor in PDB entry 7a3m for this water is 9.1 Å^2^ similar to those of the methyl carbons of the side-chains of residues 49 and 57 (9.5 Å^2^ and 9.8 Å^2^ for the delta carbons in Leu-49, and 8.9 Å^2^ and 9.3 Å^2^ for the gamma carbons in Val-57). The other water molecules in this region are more accessible to the bulk solvent but still have low B-factors (11 Å^2^ – 12 Å^2^). The water molecules near the end of the β-strands where we observe the most pronounced slow dynamics on backbone N^H^ have higher B-factors at 13.6 Å^2^, 17.1 Å^2^ and 18.5 Å^2^ for waters 144, 125 and 134. As these water molecules and the residues that undergo slow conformational exchange interact, we suggest that the process giving rise to this exchange is the binding and release of one or more of the water molecules to bulk water. The negative value for Δ*C*p that we determined for the slow exchange process would indeed be compatible with a process where bound water molecules are released to bulk solvent^27^. We also speculate that the release of water molecules and the direct contact between water 150 and the side-chains of residues 49 and 57 could be the mechanism for the differences in stability of minor state that we observe for the four CI2 variants studied in this work.

In conclusion, we have demonstrated that CI2 undergoes a slow conformational exchange process that may involve the release of water molecules bound to specific sites on the protein. The small size of CI2 has always made this protein an attractive model system in protein science. The slow dynamic process that we have characterized here could make CI2 also of interest for experimental analysis of the role of long-lived water molecules in protein structures and for benchmarking long milli-second MD simulations.

## Methods

### Protein production and sample preparation

WT (UniProt: P01053, residues 22-84 with an additional N-terminal Met) and mutated variants of CI2 (L49I, I57V, and L49I/I57V) were expressed using the previously described DNA constructs^9^. The proteins were produced in *E. coli* BL21(DE3) carrying the expression plasmids. The bacterial cultures were grown at 37 °C in M9 minimal medium with ^15^N-amoniumchloride as the sole nitrogen source. The temperature was lowered to 20 °C and the expression was induced with 0.4 mM isopropyl b-D-1-thiogalactopyranoside at OD_600_ = 0.6. After overnight growth cells were harvested by centrifugation at 5000 g and the pellet resuspended in 25 mM Tris HCl, 1 mM EDTA, pH 8. Cell lysis was performed by two freeze-thaw cycles. Insoluble material was separated from soluble protein by centrifugation at 30000 g for 20 minutes at 5 °C. Nucleic acids were precipitated with 2% polyethylene imine and removed by centrifugation at 20,000 g for 15 min at 5°C. CI2 was precipitated by adding ammonium sulfate to 70% saturation and centrifuging at 20,000 g for 15 min at 5 °C. The ammonium sulfate precipitate was resuspended in 10 mM ammonium bicarbonate, and pure CI2 was obtained by SEC on a Superdex 75 26/600 in 10 mM ammonium bicarbonate followed by lyophilization.

### NMR Experiments

All NMR experiments were performed with samples containing 1.5 mM CI2 in 50 mM MES, pH 6.26, 5% D_2_O, and 0.02% NaN_3_. Series of ^15^N relaxation dispersion experiments were recorded on Bruker Avance HDIII 600 MHz and 750 MHz spectrometers using the single-train CW-CPMG pulse sequence^28^. At both static fields, dispersion profiles were sampled in triplicates using 9 CPMG field strengths, ν_CPMG_, from 80 to 1120 Hz. For WT CI2 all relaxation experiments were conducted at 1, 5, 10, 15, 20, and 25 °C. In case of the mutated CI2 variants the experiments were only conducted at 5 °C.

### Data analysis

All experiments were processed with nmrPipe^29^. Peak intensities were quantified by summing the data in a 3×3 point window centered on the cross-peak. The dispersion profiles were analyzed with the software *relax*^30^. At each individual temperature except 25 °C, data for each residue were fitted to a constant value and to a numerical solution to the Bloch-McConnell equations for a two-state process^31^. At 25 °C, the conformational exchange in CI2 is fast, and therefore the data were fitted to the expression of Luz and Meiboom^32^. In all cases, data sets with significant relaxation dispersion (AIC) and a dispersion step larger than 2 s^-1^ at least at one static field were included in global fits for each temperature and for each variant.

The temperature variation of Δ*G* for the conformational exchange process extracted from the relaxation dispersion data were fitted to^33^:

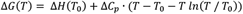

where *T*_0_ is the temperature where Δ*G* is maximal and where Δ*S* for the process is 0. Δ*C*_p_ is assumed to be independent of temperature.

## Supplementary Material

Supplementary material is available including nine figures with relaxation dispersion profiles for all residues at all conditions (Figure S1-S9), one table with a list of structures of CI2 (Table S1) and supplementary references.

## Acknowledgements

We wish to thank Dr. Johan G. Olsen for insightful help with analysis of electron density maps of CI2. This work was supported by the following grants: Carlsberg Foundation (postdoc grant no. CF17-0491 to Y.G.); the Lundbeck Foundation to the BRAINSTRUC structural biology initiative (155-2015-2666, to K.L.-L.); Novo Nordisk Foundation to the NMR infrastructure facility, cOpenNMR, (grant no. NNF18OC0032996). We also acknowledge access to the Biocomputing Core Facility at the Department of Biology, University of Copenhagen.

## Notes

### Competing Interest Statement

The authors have declared no competing interest.

