## Supplementary Material for "Slow conformational changes in the rigid and highly stable chymotrypsin inhibitor 2"

[1]Present address: Division of Biophysical Chemistry, Center for Molecular Protein Science, Department of Chemistry, Lund University, Lund, Sweden

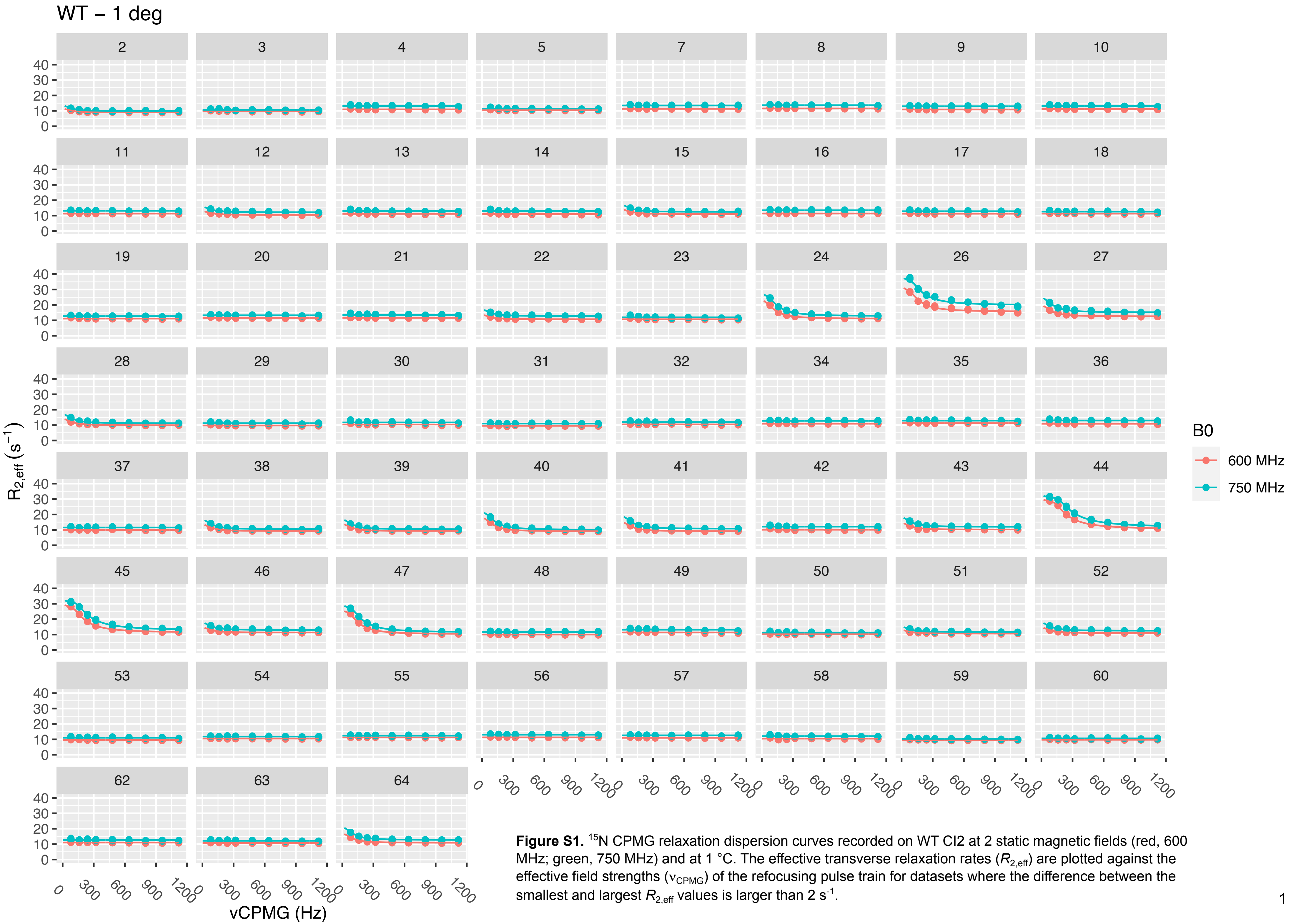


**Figure S1.** [15]N CPMG relaxation dispersion curves recorded on WT CI2 at 2 static magnetic fields (red, 600 MHz; green, 750 MHz) and at 1 °C. The effective transverse relaxation rates ($R_{2,\mathrm{eff}}$) are plotted against the effective field strengths ($\nu_{\mathrm{CPMG}}$) of the refocusing pulse train for datasets where the difference between the smallest and largest $R_{2,\mathrm{eff}}$ values is larger than 2 $s^{-1}$.

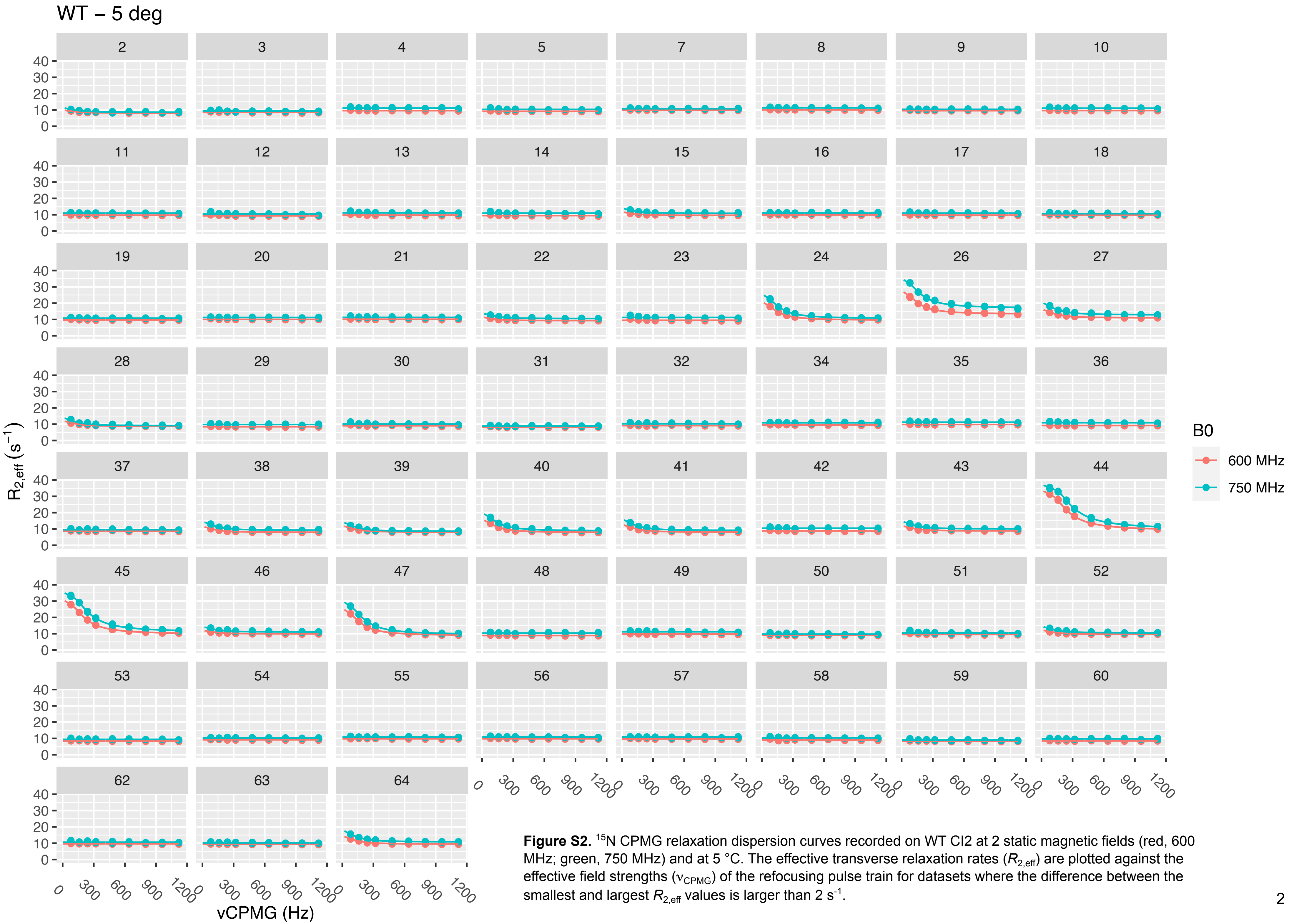


**Figure S2.** [15]N CPMG relaxation dispersion curves recorded on WT CI2 at 2 static magnetic fields (red, 600 MHz; green, 750 MHz) and at 5 °C. The effective transverse relaxation rates ($R_{2,eff}$) are plotted against the effective field strengths ($\nu_{CPMG}$) of the refocusing pulse train for datasets where the difference between the smallest and largest $R_{2,eff}$ values is larger than 2 $s^{-1}$.

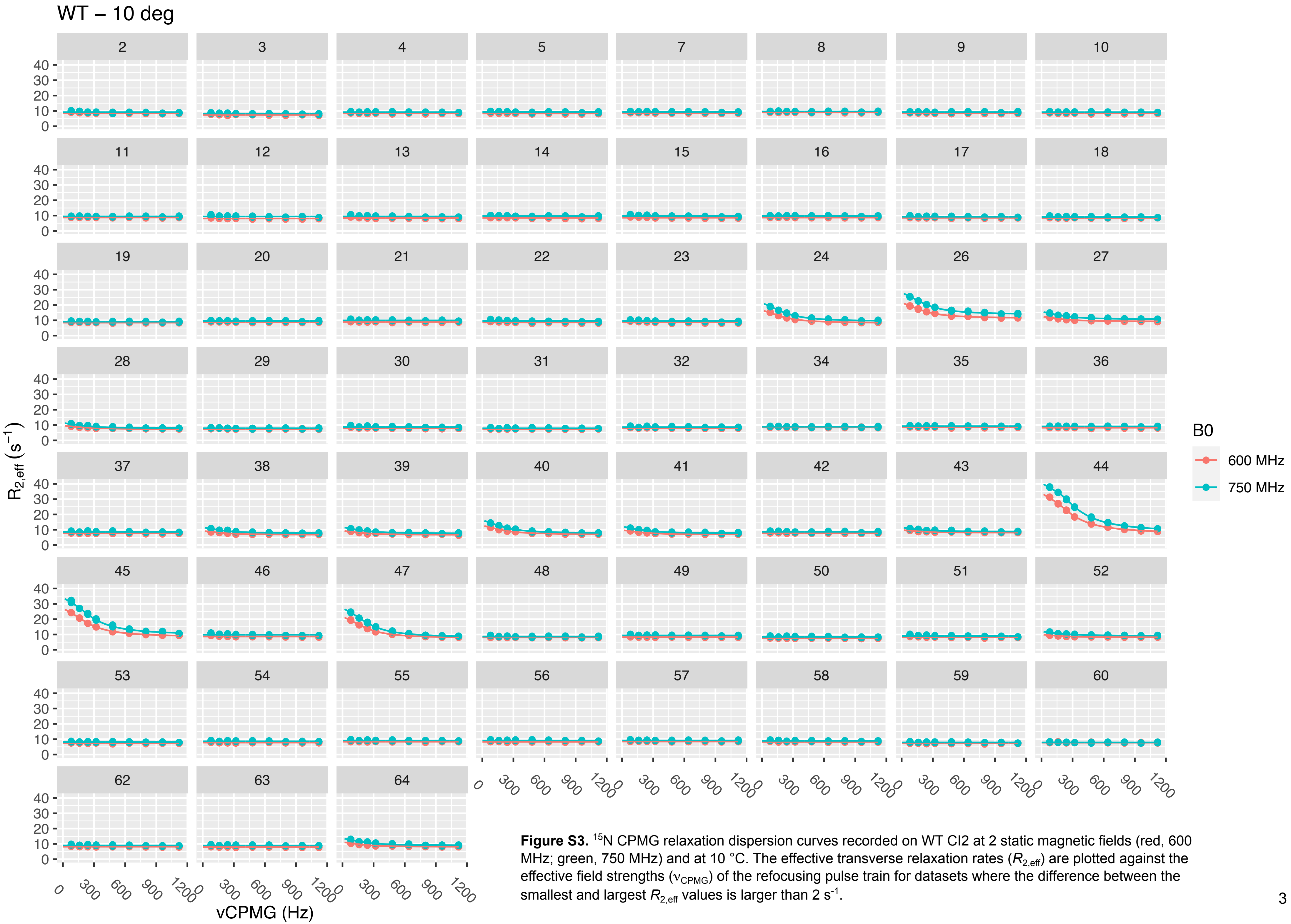


**Figure S3.** $^{15}N$ CPMG relaxation dispersion curves recorded on WT CI2 at 2 static magnetic fields (red, 600 MHz; green, 750 MHz) and at 10 °C. The effective transverse relaxation rates ($R_{2,eff}$) are plotted against the effective field strengths ($\nu_{CPMG}$) of the refocusing pulse train for datasets where the difference between the smallest and largest $R_{2,eff}$ values is larger than 2 $s^{-1}$.

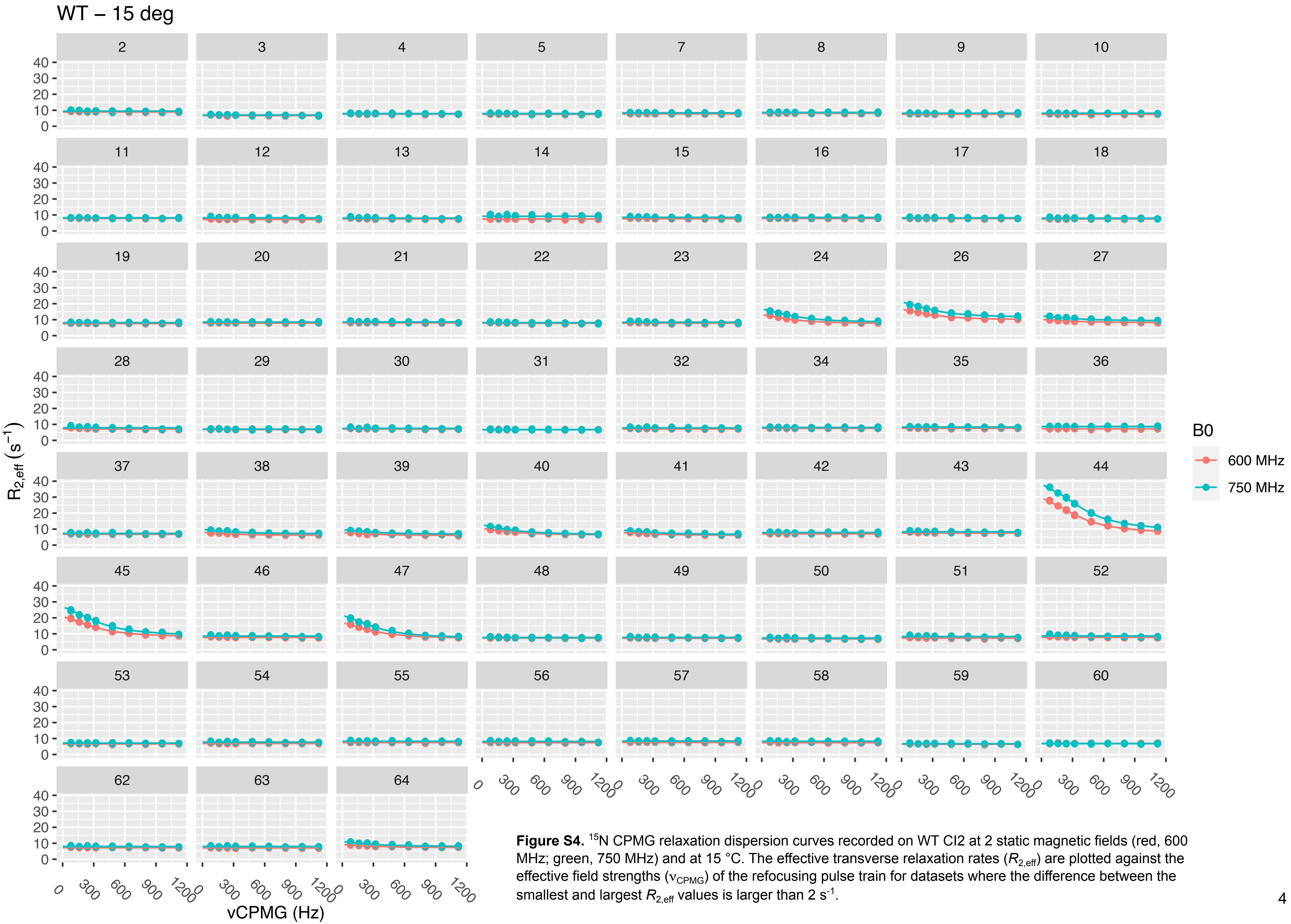


**Figure S4.** [15]N CPMG relaxation dispersion curves recorded on WT CI2 at 2 static magnetic fields (red, 600 MHz; green, 750 MHz) and at 15 °C. The effective transverse relaxation rates ($R_{2,\text{eff}}$) are plotted against the effective field strengths ($\nu_{CPMG}$) of the refocusing pulse train for datasets where the difference between the smallest and largest $R_{2,\text{eff}}$ values is larger than 2 $s^{-1}$.

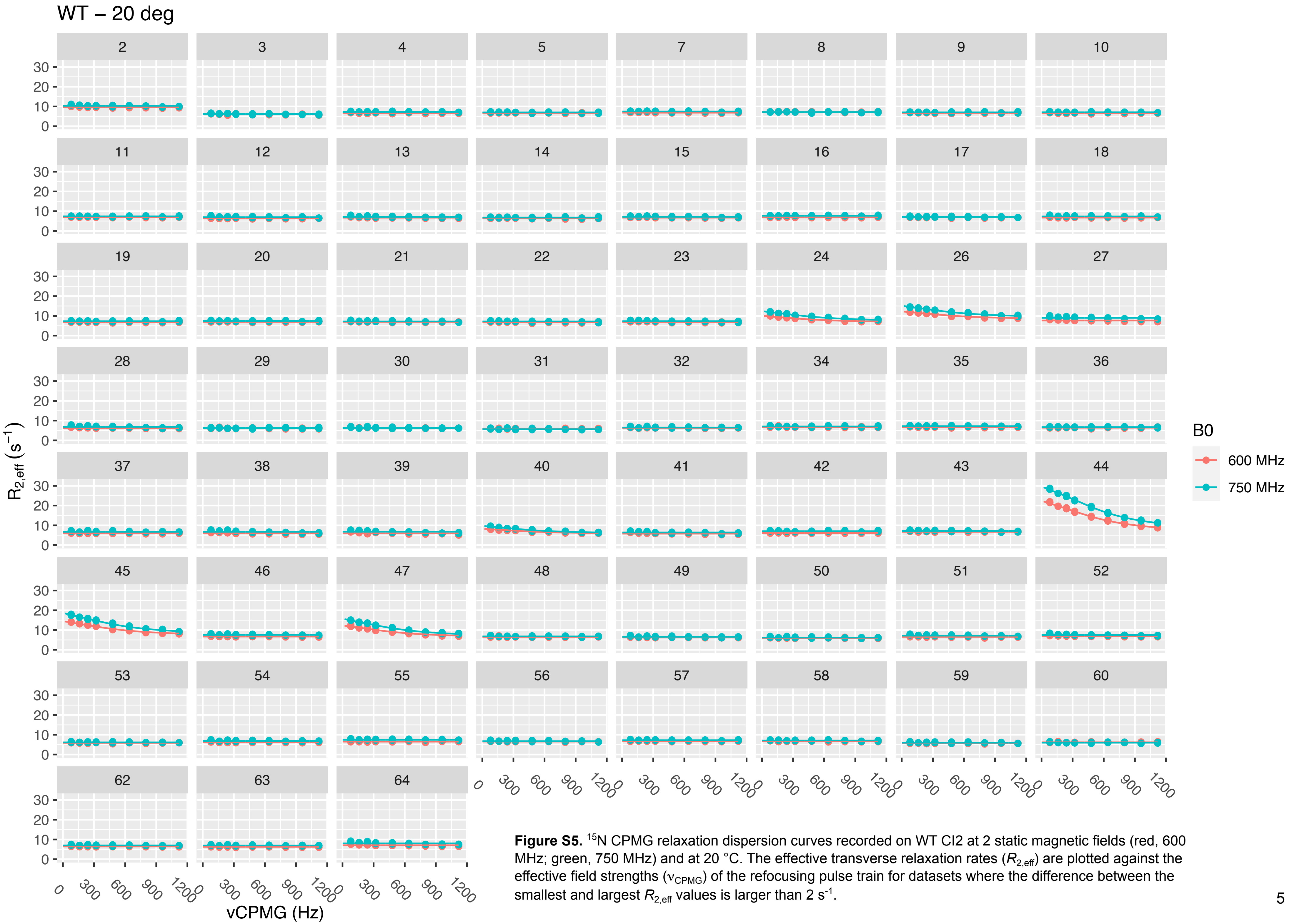


**Figure S5.** $^{15}$N CPMG relaxation dispersion curves recorded on WT CI2 at 2 static magnetic fields (red, 600 MHz; green, 750 MHz) and at 20 °C. The effective transverse relaxation rates ($R_{2,\mathrm{eff}}$) are plotted against the effective field strengths ($\nu_{\mathrm{CPMG}}$) of the refocusing pulse train for datasets where the difference between the smallest and largest $R_{2,\mathrm{eff}}$ values is larger than 2 s$^{-1}$.

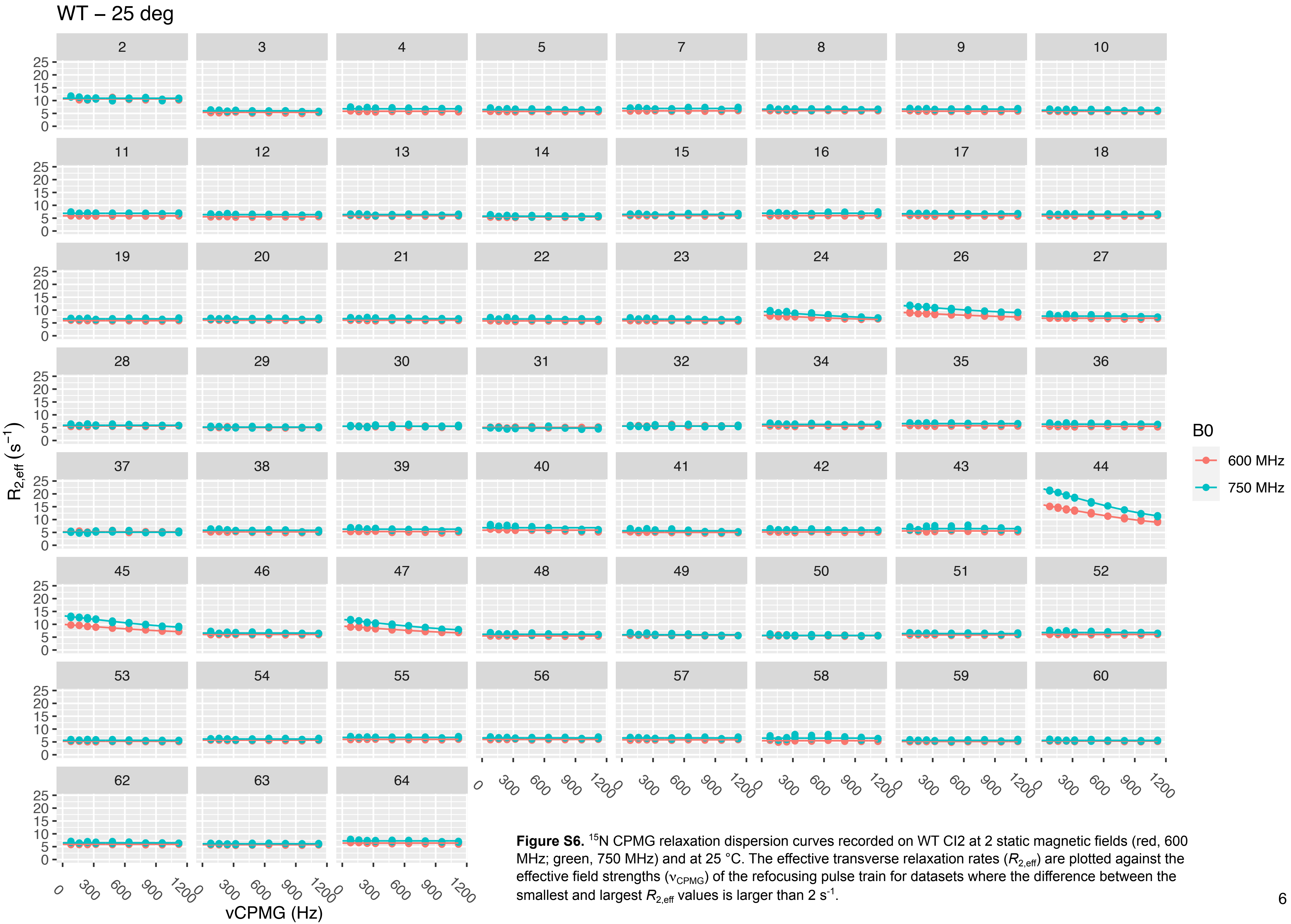


**Figure S6.** $^{15}$N CPMG relaxation dispersion curves recorded on WT CI2 at 2 static magnetic fields (red, 600 MHz; green, 750 MHz) and at 25 °C. The effective transverse relaxation rates ($R_{2,eff}$) are plotted against the effective field strengths ($\nu_{CPMG}$) of the refocusing pulse train for datasets where the difference between the smallest and largest $R_{2,eff}$ values is larger than 2 $s^{-1}$.

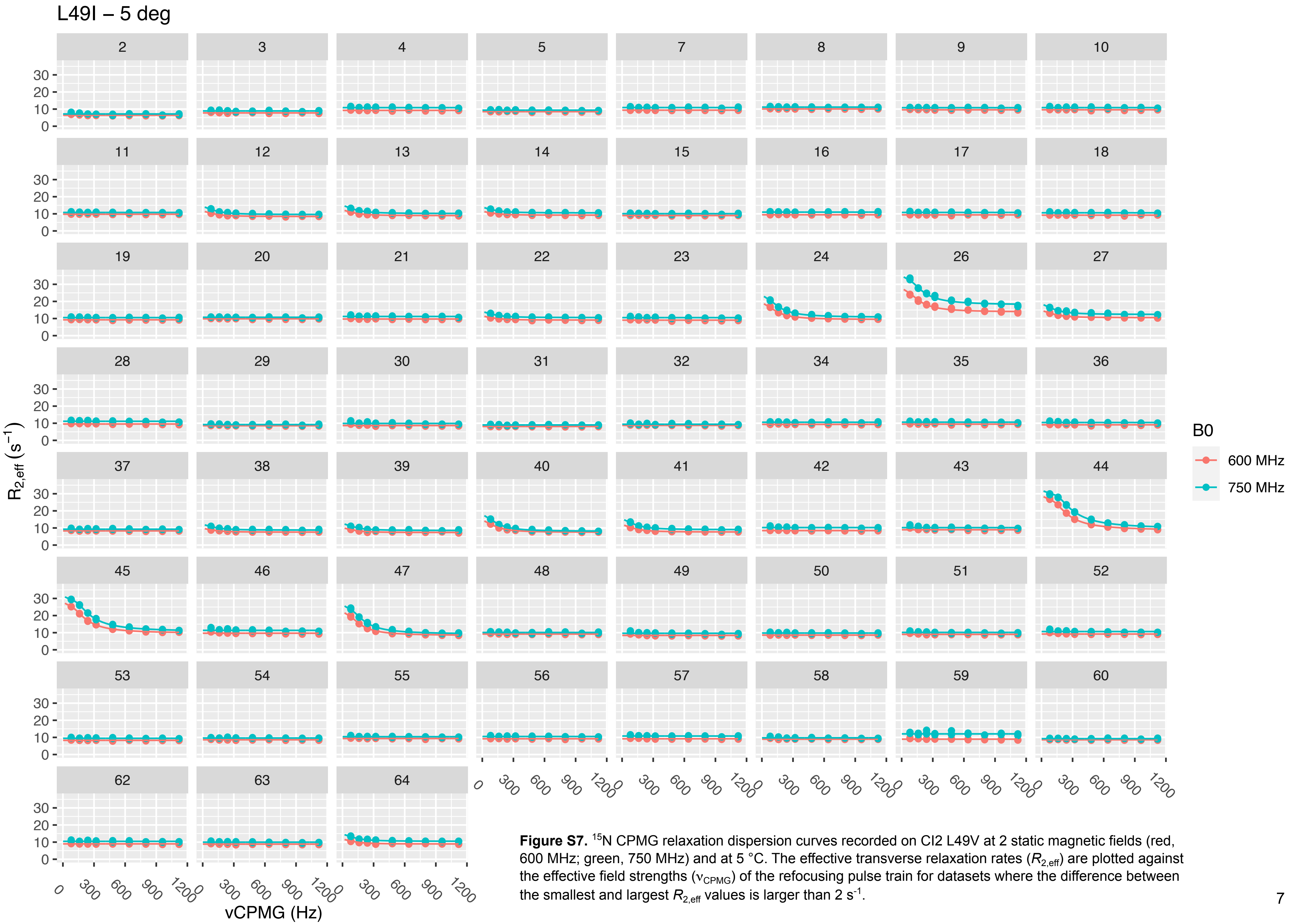


**Figure S7.** [15]N CPMG relaxation dispersion curves recorded on CI2 L49V at 2 static magnetic fields (red, 600 MHz; green, 750 MHz) and at 5 °C. The effective transverse relaxation rates ($R_{2,\mathrm{eff}}$) are plotted against the effective field strengths ($\nu_{\mathrm{CPMG}}$) of the refocusing pulse train for datasets where the difference between the smallest and largest $R_{2,\mathrm{eff}}$ values is larger than 2 $s^{-1}$.

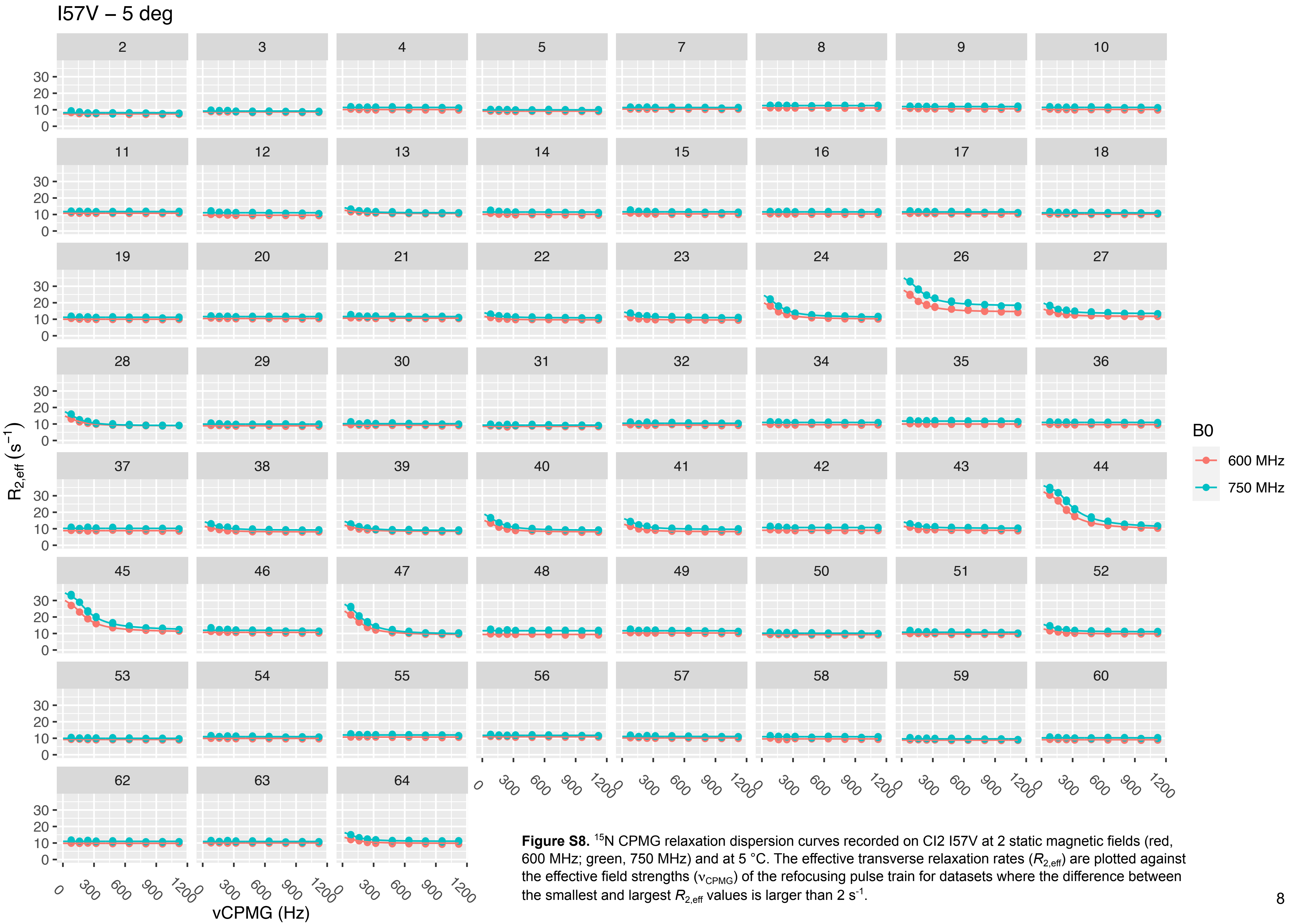


**Figure S8.** [15]N CPMG relaxation dispersion curves recorded on CI2 I57V at 2 static magnetic fields (red, 600 MHz; green, 750 MHz) and at 5 °C. The effective transverse relaxation rates ($R_{2,eff}$) are plotted against the effective field strengths ($\nu_{CPMG}$) of the refocusing pulse train for datasets where the difference between the smallest and largest $R_{2,eff}$ values is larger than 2 $s^{-1}$.

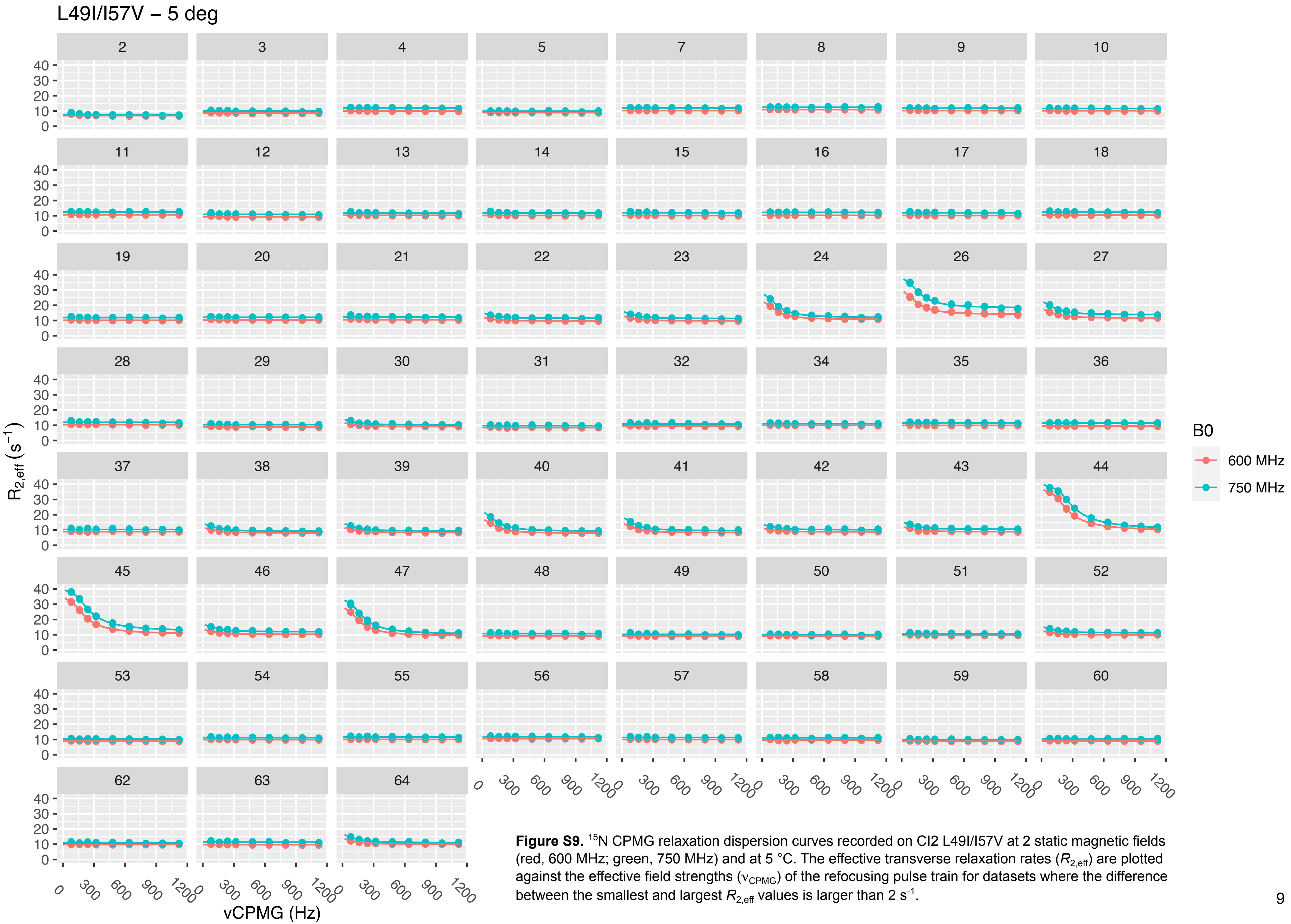


**Figure S9.** $^{15}N$ CPMG relaxation dispersion curves recorded on CI2 L49I/I57V at 2 static magnetic fields (red, 600 MHz; green, 750 MHz) and at 5 °C. The effective transverse relaxation rates ($R_{2,\mathrm{eff}}$) are plotted against the effective field strengths ($\nu_{\mathrm{CPMG}}$) of the refocusing pulse train for datasets where the difference between the smallest and largest $R_{2,\mathrm{eff}}$ values is larger than 2 $s^{-1}$.

**Table S1.** Well-defined water molecules*

| PDB entry | Six waters | Seven waters | Eight waters | Reference |
|---|---|---|---|---|
| 1COA | x | | | (Jackson et al, 1993)[1] |
| 1LW6 | | x | | (Radisky and Koshland, 2002)[2] |
| 1TM1 | | x | | (Radisky et al, 2004)[3] |
| 1TM3 | | x | | (Radisky et al, 2004)[3] |
| 1TM4 | | | x | (Radisky et al, 2004)[3] |
| 1TM5 | | | x | (Radisky et al, 2004)[3] |
| 1TM7 | | | x | (Radisky et al, 2004)[3] |
| 1TMG | | x | | (Radisky et al, 2004)[3] |
| 1TO1 | | x | | (Radisky et al, 2004)[3] |
| 1Y1K | | | x | (Radisky et al, 2005)[4] |
| 1Y33 | | x | | (Radisky et al, 2005)[4] |
| 1Y34 | | | x | (Radisky et al, 2005)[4] |
| 1Y3B | | x | | (Radisky et al, 2005)[4] |
| 1Y3C | | x | | (Radisky et al, 2005)[4] |
| 1Y3D | | x | | (Radisky et al, 2005)[4] |
| 1Y3F | | | x | (Radisky et al, 2005)[4] |
| 1Y48 | x | | | (Radisky et al, 2005)[4] |
| 1Y4A | x | | | (Radisky et al, 2005)[4] |
| 1YPA | x | | | (Harpaz et al, 1994)[5] |
| 1YPB | x | | | (Harpaz et al, 1994)[5] |
| 1YPC | x | | | (Harpaz et al, 1994)[5] |
| 2CI2 | | x | | (McPhalen and James, 1987)[6] |
| 3CI2 | --- | No waters – NMR. | --- | (Ludvigsen et al, 1991)[7] |
| 5FBZ | | x | | (Dohnalek et al, 2016)[8] |
| 5FFN | | x | | (Dohnalek et al, 2016)[8] |
| 6QIY | | | x | (Campos et al, 2019)[9] |
| 7A1H | | | x | (Hamborg et al, 2021)[10] |
| 7A3M | | | x | (Hamborg et al, 2021)[10] |
| 7AOK | | | x | (Hamborg et al, 2021)[10] |
| 7AON | | | x | (Hamborg et al, 2021)[10] |

*Number of water molecules appearing at positions in the structure of CI2 that are similar to those shown in Figure 5C.
